# Spatial and temporal representations are organized along a stable coding gradient in the medial entorhinal cortex

**DOI:** 10.64898/2026.07.09.737618

**Authors:** Hyun-Woo Lee, John C. Bowler, James G. Heys

## Abstract

The medial entorhinal cortex (MEC) has classically been viewed as a spatial coding region, but growing evidence indicates that it also contains temporal representations. However, how spatial and temporal codes are organized within the MEC remains unclear. Here we recorded MEC neurons with high-density silicon probes while mice performed a MEC-dependent timing task and a virtual reality spatial navigation task in a similar head-fixed setup. We found that spatial and temporal representations were partially overlapping but systematically biased across the MEC population. Grid cells and non-grid cells with strong spatial tuning were less likely to show reliable time-locked activity during the timing task. In contrast, neurons with weaker spatial tuning more flexibly shifted their coding scheme to temporal coding during timing task. Moreover, spatial tuning strength and its negative relationship with temporal tuning were preserved in a distinct open field environment, indicating that the coding preferences of individual neurons are constrained by a stable network-level organization. Together, these findings suggest MEC is organized along a coding gradient, ranging from dedicated stable spatial coding neurons to more flexible spatial or temporal coding neurons which represent information according to cognitive demands.

## Introduction

The medial temporal lobe (MTL) plays a critical role in encoding episodic memory by integrating what, where, and when information^1–5^. Among these dimensions, spatial coding in the MTL has received extensive attention since the discovery of hippocampal place cells^6^, leading to the identification of neurons tuned to various spatial features, including grid cells, boundary vector cells, and head-direction cells in the medial entorhinal cortex (MEC)^7–10^. These discoveries have advanced our understanding of how spatial information is represented and computed within MTL circuits and reinforced the view of MEC and hippocampus as spatial coding regions. More recently, temporal coding has been observed across multiple MTL regions. Several studies including ours have reported that neurons in the hippocampus and MEC fire at specific moments during seconds-long interval timing tasks, so-called “time cells,”^11–15^ whereas LEC exhibits slower dynamics, as its population activity drift over minutes-long epochs^16,17^. These findings establish that the MTL contains neural representations of both space and time, two fundamental components of episodic memory. However, how these representations are organized within the same neural circuit remains unclear.

MEC is particularly well positioned to address this question because it provides a major cortical input to the hippocampus^18–20^ and contains both highly structured spatial representations and activity organized by elapsed time. Therefore, a central question is what organizational principles enable MEC to support both forms of coding. At one extreme, stable circuit architecture may impose persistent, cell-specific coding biases that are preserved across behavioral contexts. At the other, coding may be dynamically reconfigured through changes in network state and task-dependent recruitment of circuit inputs, allowing neurons to flexibly represent behaviorally relevant information. Alternatively, these extremes may define the ends of a continuum of coding preferences across the MEC population. Determining how spatial and temporal coding is distributed across individual neurons can therefore provide insight into the circuit organization of MEC.

Previous studies have identified spatial and temporal coding under behavioral conditions that differed in task demands or movement state, limiting direct comparison of how these coding properties are organized in MEC.^13,14^ Resolving these alternatives requires comparing spatial and temporal coding within the same MEC population under conditions in which behavior is guided primarily by spatial position in one context and elapsed time in another while minimizing other differences in behavioral state and experimental setup. Tracking the same neurons across tasks makes it possible to determine whether individual cells retain stable functional roles or reorganize as behavioral demands change.

Here, we addressed this gap by recording MEC neurons using high-density silicon probes while mice performed a MEC-dependent interval timing task and, in the same head-fixed setup, a virtual reality navigation task. This experimental design allowed us to directly compare spatial and temporal coding in the same neurons across behavioral contexts while keeping overall movement demands and gross motor behavior broadly comparable between tasks. We found that spatial and temporal coding in MEC were partially overlapping but systematically biased. Grid cells and cells with strong spatial tuning were less likely to exhibit reliable time-locked activity, and the grid cell population preserved distance-tuned coding during the timing task. In contrast, neurons with weak spatial tuning were more likely to switch to temporal coding during the timing task. Moreover, spatial tuning strength was maintained between open-field foraging and the VR task. Together, these findings suggest that the MEC network is organized along a stable coding gradient across contexts, ranging from highly specialized spatial coding neurons to more multiplexed neurons that may combine temporal and spatial information.

## Results

### Behavioral design for comparing temporal and spatial coding in MEC

To directly compare MEC activity across temporal and spatial tasks, we established a behavioral paradigm that allowed mice to perform both tasks within a similar head-fixed setup (**Fig. 1a**). The training rig was equipped with an odor delivery spout for the timing task and surrounding monitors for virtual reality (VR)–based spatial navigation task, so that task switching can be performed by toggling the odor system and monitors either on or off. Mice (n=7) were first trained on a timing task—the temporal delayed nonmatch to sample (tDNMS) task (**Fig. 1b**)—which we developed in the previous study and used to demonstrate that the MEC is necessary for the learning of this task^15^. In each trial, a pair of odors was presented for one of three combinations of durations: long-short (L-S), short–long (S-L), and short–short (S-S). For L-S and S-L trials, mice were required to lick at the offset of the second odor to receive a water reward, whereas in S-S trials, they were expected to withhold licking as no reward is given. Thus, successful performance required the animals to accurately estimate odor durations and associate it with appropriate responses. After training, mice showed selective licking for the L-S and S-L trials (**Fig. 1c**), reaching an average performance of 80.3% (n=7 sessions, L-S: 74.6% ± 4.4, S-L: 90.6% ± 3.7, S-S: 75.9% ± 1.7; chance level = 50%; correctness is higher than chance level for all trial types- p: 0.008, Z = 28, Wilcoxon rank-sum test; **Fig. 1d**).

**Figure 1.**
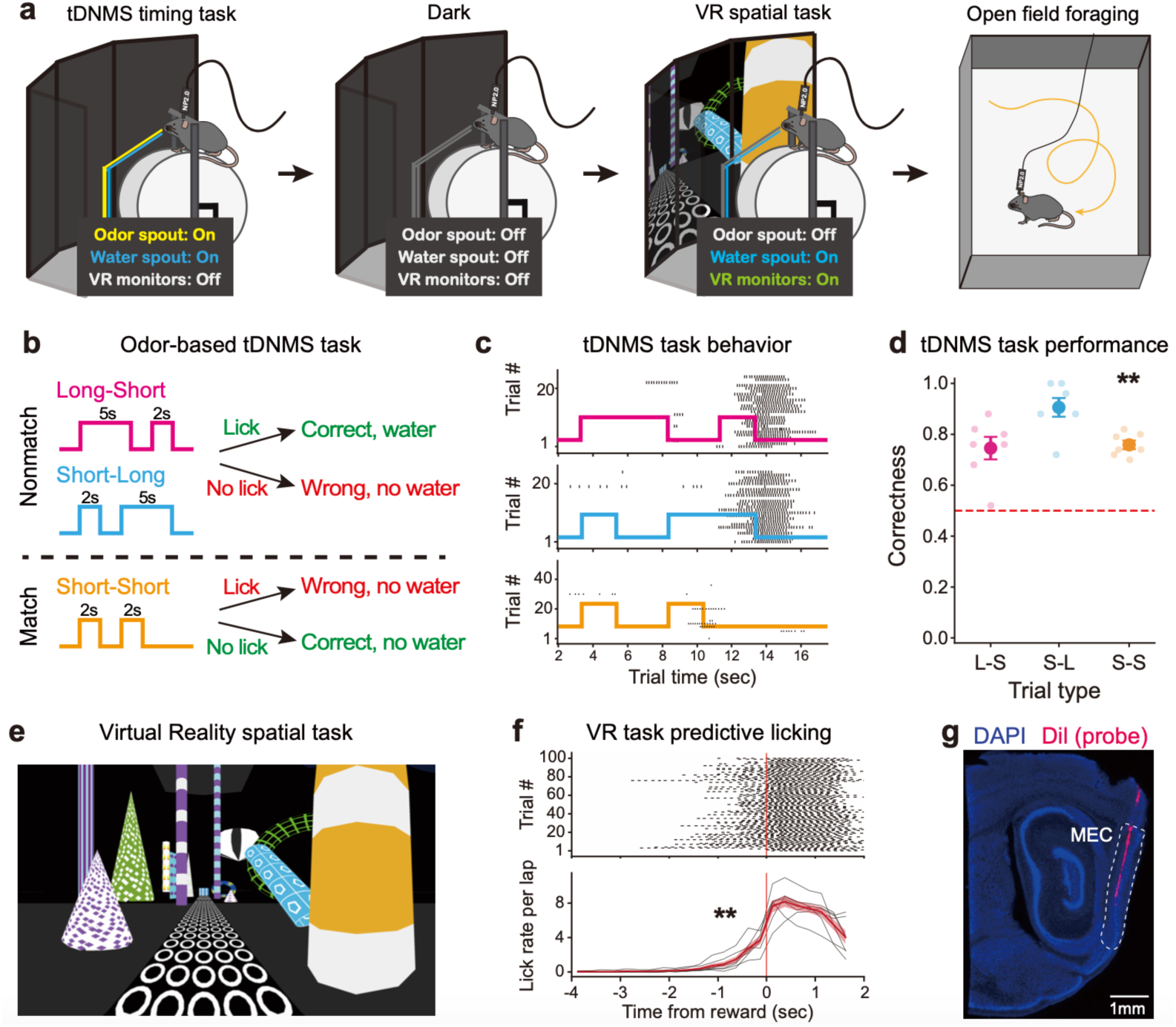
Experimental design and behavior for time and spatial tasks in head-fixed setup. **a.** An example recording sequence consisted of an odor-based timing task (tDNMS), a dark session without sensory cues or reward, a virtual reality (VR) spatial navigation task, and open field foraging. Neural activity was recorded from MEC using Neuropixels probes. **b.** Schematic of the tDNMS task. In each trial, a pair of odor durations were presented (Long-Short, Short-Long, or Short-Short), and mice were trained to discriminate the odor durations and respond accordingly. They needed to lick after the second odor to receive the reward in either Long-Short or Short-Long trials, whereas for Short-Short trials, mice needed to withhold licking. **c.** Lick raster plots from a representative session for each trial type. **d.** Behavioral performance of individual mice showing successful acquisition of the timing task. **e.** The VR environment used for the experiment from the mouse’s point of view. **f.** In the VR spatial task, mice ran along the track and received a water reward at the end of the track. Top panel shows a lick raster plot from a representative session, and the bottom panel shows average lick rate for each mouse (black lines) and the mean ± SEM across mice (red line and shaded area). All mice exhibited anticipatory licking before reward. **g.** Histology verified the Neuropixels probe track within the MEC. **p < 0.01.

Following acquisition of the timing task, mice were trained on a spatial navigation task in the same setup. In this task, mice ran a virtual linear track displayed on the monitors and received a water reward upon reaching the end of the track (**Fig. 1e**). After the reward was dispensed, they were teleported back to the starting position. Since the animals were already familiar with the head-fixed setup during timing task training, all mice adapted quickly to the VR environment and the task. To ensure mice became fully familiarized with the VR environment, we performed least four training sessions prior to recording. With training, mice exhibited stable running performance on the 3-m VR linear track, maintaining a mean running speed of 16.4 ± 2.5 cm/s across sessions. They also displayed anticipatory licking when approaching the reward zone, a hallmark behavior of spatial learning (lick rate right before the reward: 3.7 ± 0.5 per second, higher than lick rate in non-reward zone: p= 0.002, Z= 3.13, Wilcoxon rank-sum test; **Fig. 1f**).

After training, we recorded MEC neural activity using Neuropixels probes (**Fig. 1g**) while mice performed both the timing and VR tasks sequentially, with a dark session in-between each task. During the dark session, all rewards and visual or olfactory stimuli were omitted. This design allowed us to track the activity of the same MEC neurons across temporal, spatial, and spontaneous dark conditions.

### Spatial and temporal coding are partially overlapping but biased across MEC cell types

We next asked whether spatial and temporal information in MEC is represented by overlapping populations or by largely distinct subpopulations. To address this question, recorded MEC cells (n = 1,068 from 7 recording session across 7 mice, one session per mouse) were classified according to their spatial or temporal coding properties. For the spatial dimension, cells were classified into three categories: grid cells, non-grid spatial cells, and non-spatial cells. Identifying grid cells based on activity during the VR linear track was challenging, as the two-dimensional grid pattern is not readily observable in a one-dimension track. In VR, some MEC neurons exhibited multiple firing fields that repeated at the same locations across laps, resulting in highly periodic spatial autocorrelograms (**Fig. 2a**). Therefore, it is unclear how to distinguish between grid cells and non-grid spatial cells with multiple fields. To address this limitation, we leveraged neural activity recorded during the dark session. Previous studies have shown that MEC grid cells exhibit distance-dependent firing patterns when animals run on a head-fixed linear track in the absence of visual cues ^21,22^. Consistent with this, we found a subset of neurons retained distance-tuned activity during dark sessions, in the absence of any external cues such as visual stimuli or rewards (**Fig. 2a**, Cell 1∼3). To quantify this distance tuning, we computed the power spectral density (PSD) of the spike trains as a function of traveled distance, which characterizes the strength of spatial periodic firing patterns across frequencies. Cells were classified as grid cells if the PSD displayed a significant peak relative to shuffle controls (**Fig. 2a**, Cell 1∼3). Consistent with previous reports, we observed discrete jumps in the spacing of periodic firing along a gradient from dorsal to ventral MEC in grid cells identified using PSD (**Fig. 2b**) ^8,22–24^.

**Figure 2.**
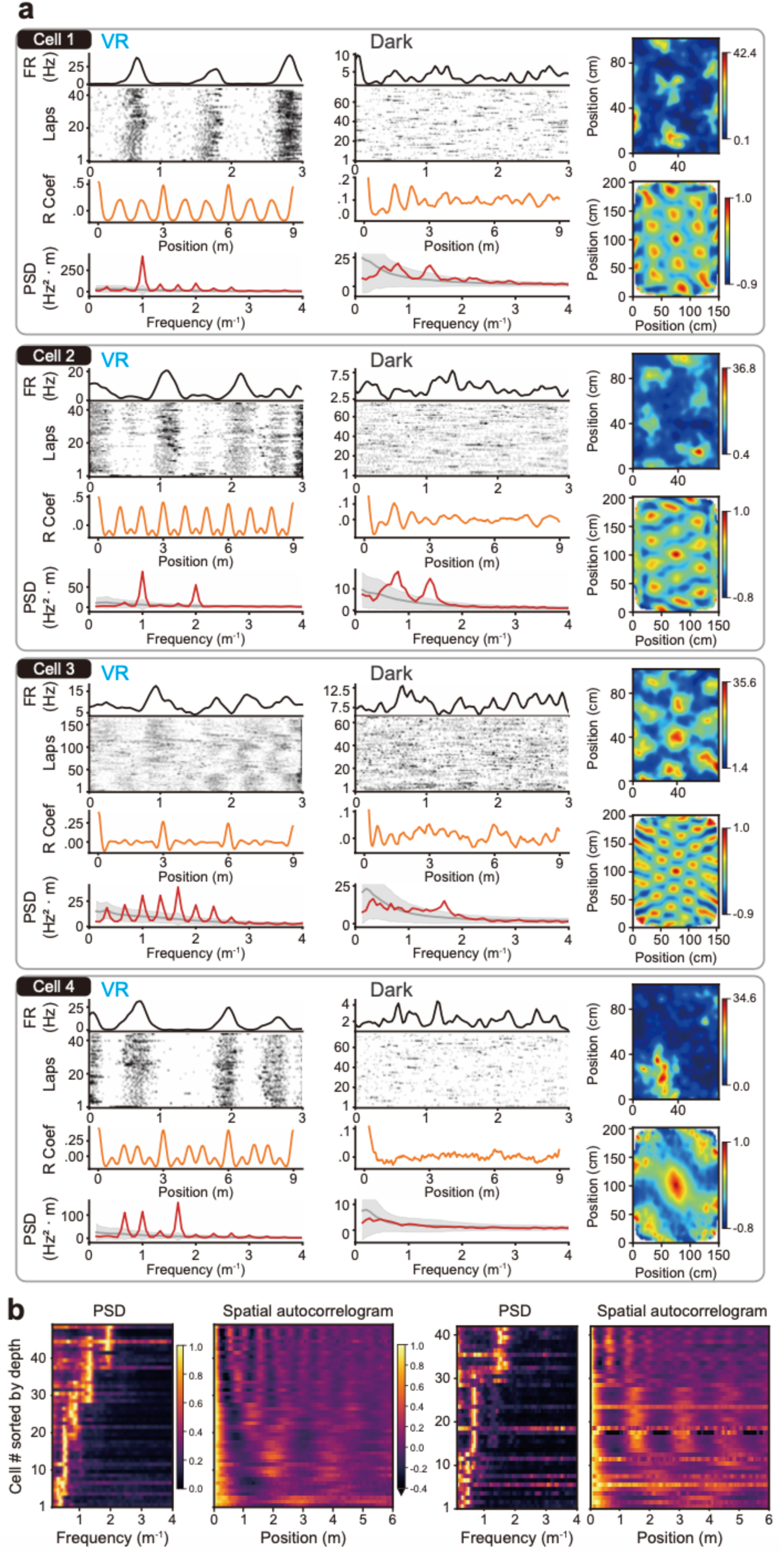
Identification of grid cells using distance-tuned activity during the Dark session. **a.** Representative activity of MEC neurons during the VR task (left), the Dark session (middle), and open field foraging task (right). The VR and Dark panels consist of an average firing rate map across laps, spike raster plots, spatial autocorrelogram, and power spectral density (PSD) of firing as a function of traveled distance. In the PSD plot, red line represents the observed PSD, and the PSD distribution of shuffled data is shown in gray (mean ± 5 SD). The open field foraging panels show both rate maps and corresponding 2-D spatial autocorrelogram. Cells with PSD peaks exceeding the mean + 5 SD of shuffled data during the Dark were classified as grid cells. **b.** PSD and corresponding 1-D spatial autocorrelograms of identified grid cells sorted by anatomical depth from two representative recording sessions (left and right). Spatial periodic scale increased discretely along the dorsoventral axis, consistent with the known properties of 2D grid cells.

To further validate this classification, we recorded neural activity in a separate set of sessions consisting of the timing task, a dark period, the spatial task, and a two-dimensional open field foraging task with scattered food (n = 825 cells from 4 recording sessions across 4 mice, one session per mouse). The overlap between 2-D grid cells and 1-D distance-tuned cells was significantly greater than expected by chance (**Fig. 2a**; 2-D grid cells n=110, 1-D distance-tuned cells n=181, overlap between them = 60; odds ratio = 5.89, p = 4.17 × 10^-16^, Fisher’s exact test), indicating a strong association between distance tuning in the absence of external cues and canonical grid cell properties observed in 2-D space. Building on this overlap, we compared the spatial periodicity of these neurons across dimensions. For cells identified as both 2-D grid cells and 1-D distance-tuned cells, the grid spacing measured in the 2-D open field was positively correlated with the spatial period observed in the 1-D condition (Pearson correlation r = 0.38, p = 0.002), indicating a preserved relationship in their spatial tuning across environments. Notably, the spacing measured in 1-D was significantly larger than that observed in 2-D (2-D spacing mean: 45 cm, 1-D spacing mean: 72 cm; Z = 12.0, p = 2.51 × 10^-11^, Wilcoxon signed-rank test), consistent with the previous findings ^25,26^. These results support the use of distance-dependent firing in dark session as a reliable proxy for identifying grid cells.

Cells that exhibited spatially selective activity during the VR session but did not meet grid cell criteria were classified as non-grid spatial cells, and the remaining cells with weak or no spatial modulation were categorized as non-spatial cells (grid cell n = 222, spatial cell n = 294, non-spatial cell n = 334).

In the tDNMS task, mice must estimate odor duration to guide their behavioral response. We therefore classified neurons based on whether they encoded temporal information during the tDNMS task. Cells exhibiting reliable temporally structured activity across trials were classified as time cells, whereas cells lacking such trial-by-trial reliability were categorized as non–time cells^27^ (time cell n = 279, non-time cell n = 571).

To determine the organization of spatial and temporal information within MEC, we examined the cells’ activity pattern across tasks. We observed that grid cells often lacked time-locked activity during the timing task (**Fig. 3a**, Cell 1) alongside cells with strong spatial tuning during VR **Fig. 3a**, Cell 2). Conversely, cells classified as time cells were less likely to show distance-tuned activity or strong spatial tuning (**Fig. 3a**, Cell 3).

**Figure 3.**
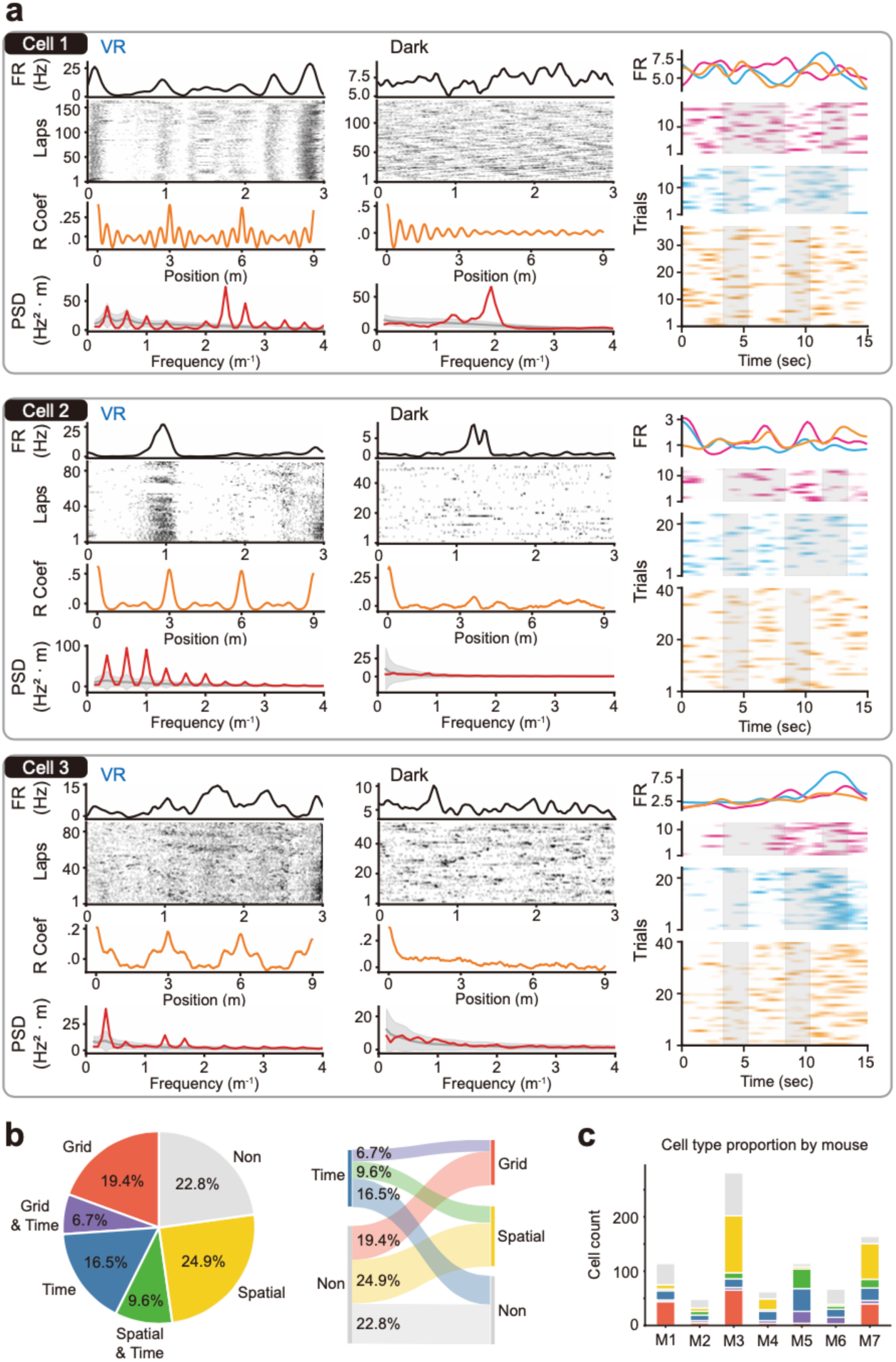
Spatial and temporal information are represented by largely segregated populations in the MEC. **a.** Representative neurons for each cell type (grid cell, spatial cell, and time cell) across the VR, Dark, and tDNMS tasks. Left and middle panels are the same as those in Figure 2a. Right panel shows activity during the tDNMS task, with average rate map (top) and trial rate maps for each trial type (bottom). Gray shading indicates odor presentation. **b.** Proportion of spatial and temporal cell types (left) and Sankey diagram showing transitions between spatial and temporal classifications (right). Grid cells and spatial cells exhibited less overlap with time cells than expected by chance. **c.** Distribution of cell types in individual mice (n=7).

This observation was consistent at the population level. The overlap between grid cells and time cells (6.7% of all neurons) was significantly smaller than expected by chance (**Fig. 3b**; p = 0.0098, Fisher’s exact test). Likewise, the overlap between non-grid spatial cells and time cells (9.6% of all neurons) was also significantly lower than expected by chance (**Fig. 3b**; p = 0.026, Fisher’s exact test). This pattern was observed consistently across a majority of mice (**Fig. 3c**). Together, these results indicate that temporal coding is not uniformly distributed across MEC spatial cell types. Instead, time cell activity is less prevalent among grid and spatial cells, suggesting a biased, partially overlapping organization of spatial and temporal coding in MEC.

### Coding flexibility varies along the strength of spatial tuning

At the population level, we observed limited overlap between spatially tuned cells and time cells. This raised the question of how coding preferences vary at the level of individual neurons: which MEC neurons flexibly change their coding according to task demands, and which neurons maintain more stable coding across tasks? We therefore examined spatial and temporal tuning strength at the single-cell level across behavioral tasks. Spatial tuning during the VR task was quantified using spatial mutual information (MI), and temporal tuning during the tDNMS task was assessed by trial-by-trial reliability, measured as the Pearson correlation coefficient across trials.

We also performed a generalized linear model (GLM) analysis to determine whether each neuron’s activity was better explained by elapsed distance or elapsed time, given that conventional tuning metrics alone cannot fully dissociate spatial and temporal coding. For example, a neuron that fires according to elapsed time during the VR task may appear spatially tuned because elapsed time and position are highly correlated. Likewise, in the tDNMS task neurons may appear temporally tuned when they encode traveled distance. For each neuron, we compared a spatial model containing elapsed distance and a temporal model containing elapsed time against a null model and quantified the improvement in model performance using the log-likelihood increase (LLHi). Negative LLHi values indicate that the corresponding model did not outperform the null model. To directly compare the relative contribution of spatial and temporal variables, we further calculated ΔLLHi (LLHi_spatial − LLHi_temporal), where positive values indicate stronger spatial coding and negative values indicate stronger temporal coding.

GLM analysis revealed a task-dependent shift in coding preference. The proportion of neurons better explained by the spatial model decreased dramatically from 99.8% during the VR task to 14.2% during the tDNMS task (n = 850, χ^2^ = 667.5, p < 0.001, McNemar’s test). In contrast, the proportion of neurons better explained by the temporal model increased from 2.0% in the VR task to 24.0% during the tDNMS task (n = 850, χ^2^ = 160.9, p < 0.001, McNemar’s test). Consistent with this shift, the distribution of ΔLLHi also shifted significantly toward negative values during the tDNMS task compared with the VR task (**Fig. 4a**; Z = 24.6, p < 0.001, Wilcoxon rank-sum test), indicating an overall increase in temporal coding preference across the population. These results indicate that coding preference across the MEC population is dynamically reorganized according to task demands and the external sensory information.

**Figure 4.**
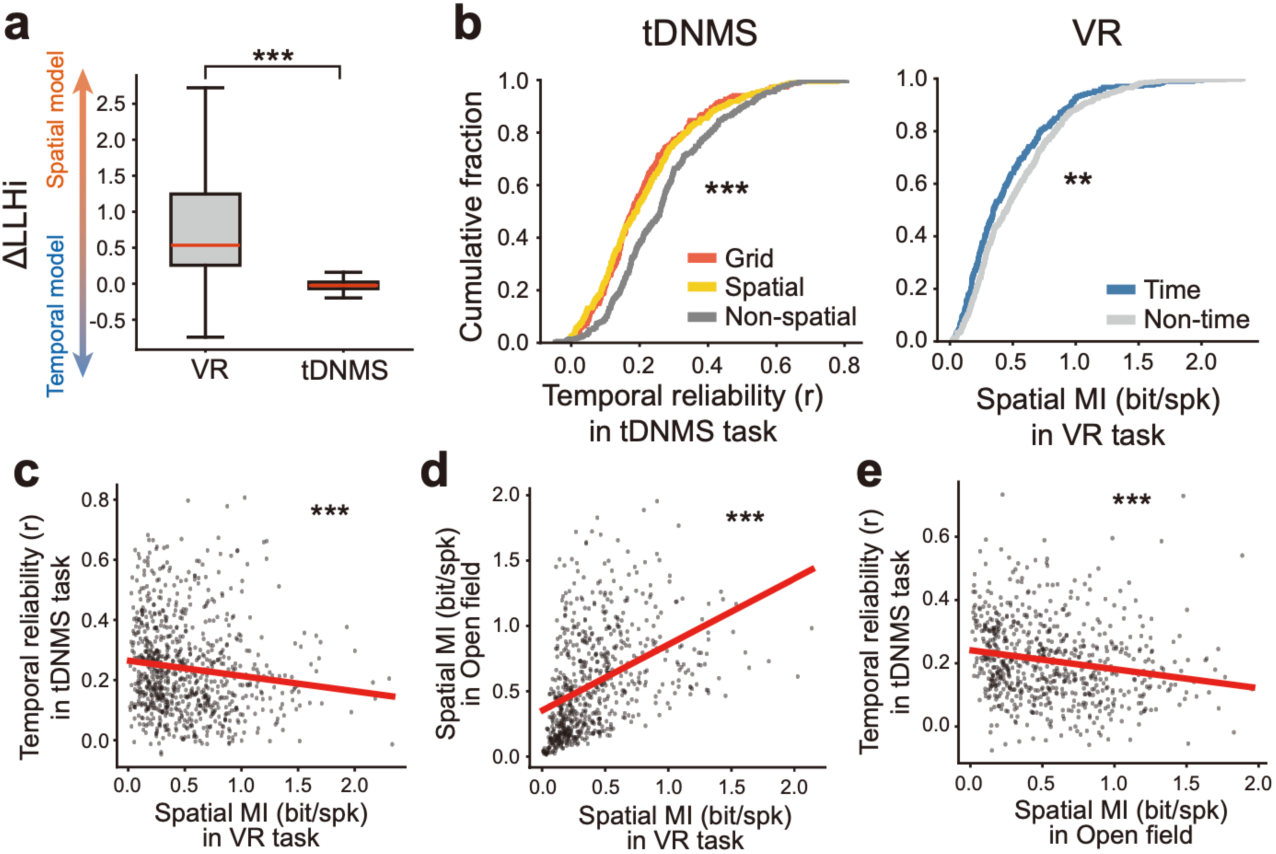
Inhomogeneous coding flexibility across the MEC population. **a.** Difference in GLM performance between spatial and temporal models (Δ*LLHi* = *LLHi_S_* − *LLHi_T_*) during the VR and tDNMS tasks. MEC neural activity was predominantly explained by position during the spatial task, and elapsed time better explained neural activity during the timing task. **b.** Spatial or temporal tuning across cell types and tasks. Left panel shows that non-spatial cells exhibited stronger temporal tuning than grid and spatial cells. Right panel shows that non-time cells exhibited stronger spatial tuning than time cells. **c.** Spatial tuning during the VR task and temporal tuning during the timing task across all MEC principal cells were negatively correlated. **d.** Spatial tuning strength of individual MEC neurons maintained across the VR task and open field foraging. **e.** Spatial tuning during the open field foraging and temporal tuning during the timing task were negatively correlated. **p < 0.01, ***p < 0.001.

Although the preceding analyses revealed a population-wide shift from spatial to temporal coding, they did not determine whether this shift occurred uniformly across MEC neurons or was constrained by a neuron’s functional identity. Because grid cells are thought to participate in a highly structured network that maintains stable relationships among cells, this network architecture could constrain their ability to acquire task-dependent temporal representations. We therefore asked which neurons were more likely to shift their coding preference across tasks. During the VR task, the activity of all cell types including grid cells, spatial cells, and non-spatial cells displayed stronger spatial tuning over temporal tuning (ΔLLHi grid cells: 1.35 ± 0.09, W = 32, p < 0.001; spatial cells: 1.19 ± 0.06, W = 63, p < 0.001, non-spatial cells: 0.34 ± 0.03, W = 2236, p < 0.001; Wilcoxon signed-rank test), and grid cells and spatial cells exhibited significantly higher spatial MI than non-spatial cells (p < 0.001; Wilcoxon rank-sum test). During the tDNMS task, however, grid cells and spatial cells exhibited significantly weaker temporal tuning than non-spatial cells (**Fig. 4b**, left; p < 0.001; Wilcoxon rank-sum test). Consistently, non-spatial cells represented the highest proportion of neurons preferentially explained by the temporal model as compared with grid and spatial cells (grid cells: 19.4%, spatial cells: 21.1%, non-spatial cells: 29.6%, χ^2^ = 9.8, p = 0.007, chi-square test of independence).

To further validate this finding from the opposite perspective, we classified neurons as time cells or non-time cells based on their activity during the tDNMS task and then examined their spatial coding in the VR task. We compared the spatial coding strength of time cells and non-time cells to determine whether temporal coding identity predicted spatial coding in a distinct behavioral context. GLM analysis showed that the activity of time cells during tDNMS task was better explained by elapsed time than traveled distance (ΔLLHi = -0.065 ± 0.009, W = 2072, p < 0.001, Wilcoxon signed-rank test), indicating that the trial-by-trial consistency of time cell activity reflected elapsed time rather than a confounding effect of traveled distance. During the VR task, time cells showed lower spatial MI than non-time cells (**Fig. 4b**, right; Z = 3.1, p = 0.002, Wilcoxon rank-sum test). Nevertheless, their activity in the VR task was still better explained by the spatial model than the temporal model (ΔLLHi = 0.79 ± 0.06, W = 1123, p < 0.001, Wilcoxon signed-rank test), indicating that their coding preference shifted toward spatial information under spatial task demands. Consistent with these results, spatial tuning measured during the VR task was significantly negatively correlated with temporal tuning measured during the tDNMS task (**Fig. 4c**; r = -0.13, p < 0.001, Pearson correlation), indicating that neurons with stronger spatial representations tended to exhibit less reliable temporal responses across tasks.

To test whether this bias in coding flexibility generalized across distinct spatial contexts, we examined sessions in which mice sequentially performed open-field foraging, tDNMS, Dark, and VR tasks (Separate sessions from the above analysis. Four sessions from four mice). We found that spatial mutual information was significantly correlated between the one-dimensional VR task and two-dimensional open field foraging (**Fig. 4d**; r = 0.43, p < 0.001, Pearson correlation), even though these tasks differed substantially in behavioral state and sensory context: VR navigation was performed head-fixed along a linear virtual track, whereas in open field foraging mice freely explored a two-dimensional arena located in a different room. Because grid cells are known to preserve grid firing pattern across environments, we repeated this analysis after excluding grid cells to determine whether this relationship was driven solely by grid cells. Spatial tuning in VR remained significantly correlated with spatial tuning in the open field (r = 0.53, p < 0.001), indicating that non-grid cell population exhibited a stable spatial coding tendency as well. We then examined the relationship between this stable spatial coding and temporal tuning during tDNMS task. Consistent with the VR results, open field spatial tuning was negatively correlated with temporal tuning (**Fig. 4e**; r = -0.19, p < 0.001), and time cells showed lower open field spatial tuning than non-time cells (Z = 4.78, p < 0.001, Wilcoxon rank-sum test).

Together, these results suggest that MEC coding flexibility is constrained by a structured organization rather than randomly assigned across population. Neurons with strong spatial tuning tended to preserve spatial modulation across environments and were less likely to exhibit temporal coding during the timing task. In contrast, neurons with weaker spatial tuning carried mixed spatial and temporal signals. Thus, MEC appears to be organized along a coding gradient, ranging from stable, strong spatial representations to more flexible multidimensional coding in which weaker spatial signals are combined with temporal information according to task demands.

### Grid cell population preserve distance-tuned activty during the timing task

Because grid cells activity is thought to reflect structured network dynamics supporting path integration, we next asked whether this network was reorganized by temporal task demands or preserved distance-tuned activity during the timing task. To examine the population dynamics of grid cells, we first identified grid modules from simultaneously recorded MEC neurons. Following a previously established approach^28^, we computed time-lagged cross-correlations of neuronal activity during the Dark session and identified modules using agglomerative clustering (**Fig. 5a**). The cross-correlation structure was highly preserved across the Dark, tDNMS, and VR tasks (**Fig. 5a**; Dark vs tDNMS: r = 0.90 ± 0.01, Dark vs VR: r = 0.45 ± 0.05), indicating that functional relationships among grid cells remained stable across behavioral contexts. Agglomerative clustering of the correlation matrix identified six grid modules across five recording sessions from three mice, and the majority of neurons within the modules were identified as grid cells (153 of 174 cells, 87.9%).

**Figure 5.**
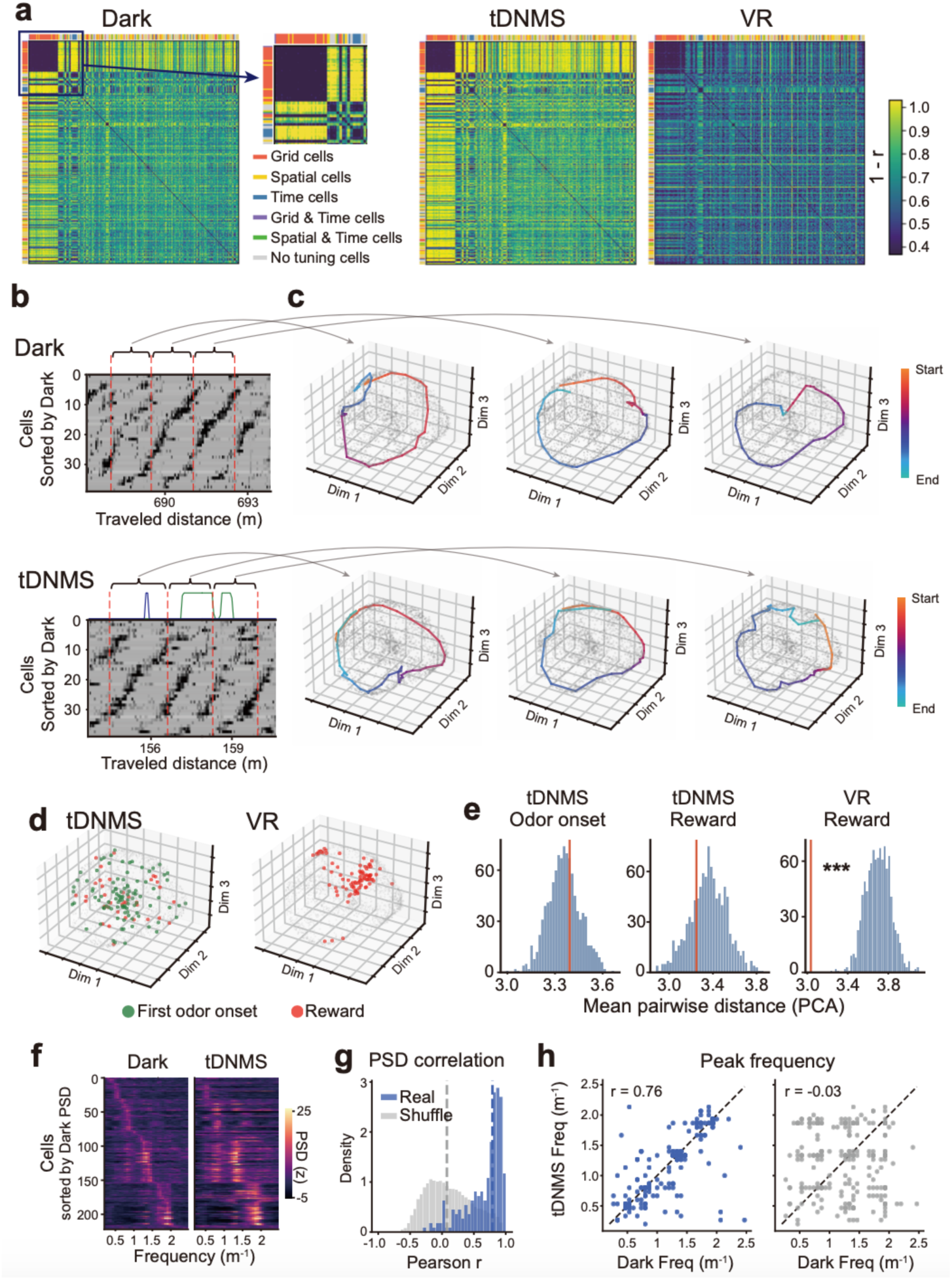
Grid cell population persists distance-tuned activity during the timing task. **a.** Time-lagged cross correlation matrices of simultaneously recorded MEC neurons during the Dark, tDNMS, and VR tasks. Neurons were sorted by agglomerative clustering of the correlation matrix during the Dark, and the same cell ordering was applied to the other tasks. The inset shows zoomed-in view of a highly correlated cluster, identified as a grid module, in the ensemble. **b.** Population activity of a representative grid module plotted as a function of traveled distance during the Dark (top) and the tDNMS task (bottom). Cells were sorted in the same order for the Dark and tDNMS tasks and showed similar sequential activity across tasks. **c.** Representative grid module activity was projected into a low-dimensional embedding. The projector was computed based on the neural activity during Dark and applied to activity during both Dark and tDNMS. Background gray dots represent population states and lines show state trajectories during the specified epochs in b. Population trajectories were similar across the two tasks. **d.** Population states corresponding to task events projected onto the same state space as in c. Green, first odor onset; red, reward delivery. **e.** Mean pairwise distances of population activity associated with task events. Population activity associated with task events formed distinct clusters during the VR task but not during the tDNMS task. Red line: observed data. Blue histogram: shuffled data. **f.** The PSDs of all grid cells were similar between the Dark and tDNMS tasks. Cells were sorted by peak frequency of PSD during Dark. **g.** Distribution of correlations between PSDs during the Dark and tDNMS tasks was significantly higher than shuffled data. **h.** Peak frequencies of PSD during the Dark and tDNMS tasks were highly correlated (left), whereas cell identity shuffled data showed no significant correlation (right). **p < 0.01, ***p < 0.001.

We next visualized ensemble activity within each module as a function of traveled distance. During the Dark session, ensemble activity exhibited a clear sequential pattern along traveled distance (**Fig. 5b**). Interestingly, applying the identical cell ordering to activity during the tDNMS task revealed a highly similar sequential organization (**Fig. 5b**), suggesting that grid cell ensemble dynamics were largely preserved across behavioral tasks. To characterize these population dynamics quantitatively, we projected Dark session ensemble activity into a low-dimensional state space using PCA followed by UMAP and projected the tDNMS activity onto the same embedding (**Fig. 5c**). Population activity during the two tasks seemed to follow highly similar trajectories in the shared state space, indicating that the dynamics of grid cell population remained largely unchanged in the timing task.

If grid-cell populations encoded task events, activity associated with specific trial events would be expected to occupy similar regions of the state space across trials. Consistent with this expectation, reward-related activity during the VR task formed clusters (**Fig. 5d, e**). In contrast, activity corresponding to trial onset and reward delivery during the tDNMS task were scattered throughout the state space (**Fig. 5d, e**). These results indicate that grid cell population dynamics were reliably aligned to task events during the spatial task but not during the timing task.

This persistence of distance-tuned activity of grid cells was also observed at the single-cell level. The PSD plots of individual grid cells showed highly similar spectral profiles between the Dark and tDNMS sessions (**Fig. 5f, g**; Z = 19.3, p < 0.001, Wilcoxon rank-sum test). Furthermore, the dominant spatial frequency of each grid cell was strongly correlated across the two conditions (**Fig. 5h**; empirical p < 0.001 compared with the shuffle distribution). The results were similar when the tDNMS session was analyzed separately during the trial phase and inter-trial intervals. Together, these findings demonstrate that grid cells activity largely maintains distance-tuned activity pattern even during the timing behavior.

## Discussion

Our results reveal a structured organization of spatial and temporal coding in MEC. By leveraging high-throughput single-unit neural recordings of the same MEC neurons across a spatial task and a temporal task in a similar setup, we found that the spatial and temporal representations were partially overlapping but systematically biased across the MEC population. Cells with strong spatial tuning were less likely to exhibit reliable time-locked activity, especially the grid cell population which displayed persistent distance-tuned activity during the timing task. In contrast, neurons with weaker spatial tuning were more likely to show temporal coding. Interestingly, this gradient of coding property ranging from strong spatial coding to multi-domain coding was largely maintained across environments.

These findings provide insight into the functional organization of the MEC network. The notable stability of each neuron’s coding profile across environments suggest that MEC neurons’ coding preferences may be determined in part by the structure of intrinsic recurrent connectivity and feedforward input. Interestingly, a similar cell-specific coding propensity has been reported in the downstream hippocampus, where individual neurons differ in their intrinsic tendency to form place fields – a property which is maintained across time and contexts^29,30^. This raises the possibility that stable coding biases could represent a general organizing principle across the entorhinal-hippocampal circuit. Within MEC, such biases could arise through distinct molecular and cellular mechanisms between cell types. For grid cells, local recurrent connectivity is thought to preserve pairwise activity patterns among grid cells within the same network module thereby maintaining grid cell firing patterns across environments^31–35^. For non-grid cells, reliable spatial firing can coexist with substantially greater sensitivity to environmental features than is observed in grid cells^36^, however the circuit and cellular mechanisms that enable stable cell-specific coding preferences remain unclear. One possibility is that differential weighting of feedforward inputs contributes to the coding gradient observed here. Neurons with strong spatial tuning may receive relatively stronger input from cortical regions conveying visuospatial information, such as the postrhinal cortex^37–39^, whereas neurons neurons with more flexible spatial–temporal coding may be more strongly influenced by nonspatial or contextual information streams associated with perirhinal and lateral entorhinal circuits^16,37,40,41^. Although this hypothesis remains to be tested, differences in local recurrent vs feedforward synaptic connectivity and input could jointly establish stable coding biases within MEC. Functionally, this organization could allow strongly spatially tuned cells to provide a stable spatial metric for hippocampal place maps, while mixed-selectivity MEC signals provide additional temporal and contextual information that can be integrated within the hippocampus. MEC may therefore constitute an intermediate computational stage in which spatial and nonspatial signals—including temporal information—are neither strictly segregated nor fully integrated, but instead are organized along a stable, cell-specific coding gradient that facilitates their downstream integration in the hippocampus.

Our findings provide a unifying framework for interpreting the distinct patterns of spatial and temporal coding reported in both Heys and Dombeck (2018) and Kraus et al. (2015). In Heys and Dombeck^14^, temporal and spatial coding were observed primarily during immobility and running, respectively, with very little overlap between the two coding populations. In our experiments, mice were allowed to run freely in all tasks – exhibiting spontaneous running during the timing task even though running was not explicitly required. By more closely matching behavioral state across spatial and temporal tasks, we observed a slightly greater overlap between spatial and temporal coding populations than was reported by Heys and Dombeck. Nevertheless, this overlap remained below chance level, indicating that the two populations were still significantly segregated, consistent with their findings. Kraus et al.^13^, by contrast, reported a high degree of overlap between spatial and temporal coding in MEC and neighboring regions during treadmill running, with many neurons encoding both traveled distance and elapsed time. The greater overlap reported by Kraus et al. may reflect at least two key differences in task design. First, the tasks differed substantially in their physical structure. In our tasks, trial onset was signaled by non-spatial cues, such as light flashes and odor presentations. In the task used by Kraus et al., however, the time interval began when the animal stepped onto a treadmill located on the center stem of the maze, making interval onset both behaviorally salient and tied to a well-defined spatial location. This task structure may have reset the population dynamics of grid cells and other MEC neurons at treadmill entry, thereby aligning traveled distance and elapsed time signals to interval onset. Second, the tasks differed in their cognitive demands. In our VR and tDNMS tasks, animals were explicitly required to use spatial and temporal information, respectively, to guide behavioral responses, whereas the treadmill task used by Kraus et al. did not impose the same dissociation between spatial and temporal demands. These distinct task demands may have more strongly recruited task-specific neurons in our experiments, resulting in a clearer separation between spatial and temporal coding populations. Thus, our results suggest that the degree of overlap between spatial and temporal coding in MEC may depend on behavioral state, task structure, and cognitive demands.

An important question raised by the present study is how neurons that encode both spatial and temporal information represent these variables when they are simultaneously relevant. Because spatial and temporal coding were examined in separate behavioral tasks, recording from the same population allowed us to identify cells that encoded information in more than one domain but did not provide direct evidence that these signals are integrated at the level of individual neurons. Instead, neurons that encoded both domains in our experiment may maintain largely independent representations of spatial and temporal information rather than integrating them into a unified code. Such neurons could multiplex spatial and temporal information, dynamically switch between representations moment to moment, or preferentially represent a single domain when both types of information are simultaneously required to aid task performance. This possibility is consistent with a recent study demonstrating that MEC neurons encode visual and auditory spatial information through both unimodality and multimodality neurons^42^. Importantly, multi-modal neurons exhibited modality-dependent representations rather than a single unified spatial map shared across sensory modalities, suggesting that overlapping coding does not necessarily imply integration. Future studies using behavioral paradigms that require the simultaneous use of spatial and temporal information will therefore be important for determining how neurons with flexible coding represent these variables within the same behavioral context. Such experiments would also more closely resemble naturalistic behavior, in which animals must integrate multiple sources of information, including space, time, and sensory cues, to guide behavior.

## Methods

### Animals

Seven adult C57BL/6J mice (Charles River Laboratory; 4 males and 3 females), aged 2-3 months at the start of experiments, were used. Mice were housed on a reverse 12 h light/ 12 h dark cycle under controlled temperature (approximately 21 °C) and humidity (25-45 %) conditions. All experimental procedures were approved and conducted in accordance with the University of Utah Institutional Animal Care and Use Committee.

### Headplate and Neuropixels probes implantation

Headplate and Neuropixels probe implantation procedures were performed as previously described^27^. Briefly, mice were anesthetized with 1–2% isoflurane. A custom titanium headplate was attached to the skull using Metabond (Parkell), and skull screws were implanted over the left hemisphere and cerebellum. The cerebellar screw was connected to a silver wire (teflon coated silver wire, 30 AWG, WPI), which served as the recording ground. For probe implantation, the tip of the Neuropixels 2.0 probe (NP2014, Imec) was sharpened using a pipette beveller (BV-10, Sutter Instrument) before mounting in a 3D-printed holder^43^. Immediately before insertion, the probe was sterilized with 99% isopropyl alcohol and then coated with a Vybrant fluorescent cell-labeling solution (Thermo Fisher) to enable post hoc verification of recording tracks. A craniotomy (∼2 × 1 mm) was performed over the right hemisphere, centered 3.3 mm lateral to bregma and exposing the anterior portion of the transverse sinus. The probe was inserted at an anterior angle of 11–13° and advanced at approximately 2 μm/s until reaching a depth of up to 3.5 mm or until a shank began to bend against the skull. The craniotomy was covered with DuraGel (Cambridge NeuroTech) and sealed with Kwik-Sil (World Precision Instruments), after which the probe assembly was fixed with Metabond. Following surgery, animals were monitored for 3 days and received daily subcutaneous injections of carprofen (5 mg/kg).

In two mice, headplate implantation was performed before behavioral training, and Neuropixels probes were implanted after animals reached criterion performance in the tDNMS task. In the remaining five mice, headplate and probe implantation were performed before behavioral training. These animals were also implanted with a second Neuropixels probe targeting the dorsal striatum; however, only recordings from the MEC are included in the present study.

### Temporal Delayed Nonmatch-to-Sample (tDNMS) task

The behavioral apparatus and training procedures for the temporal Delayed Nonmatch-to-Sample (tDNMS) task have been described previously^15^. Briefly, mice were head-fixed in a custom behavioral rig consisting of a cylindrical treadmill equipped with a rotary encoder, an odor delivery port, a lick spout for water reward delivery (∼5–6 μL), an optical lick sensor, and five LCD monitors used for virtual reality experiments. Mice were free to run on the treadmill throughout behavioral session. A 2% isoamyl acetate diluted in mineral oil (Cole-Parmer, 99+%) was used as the odor stimulus and delivered through a flow-dilution olfactometer.

Following postoperative recovery, mice were placed under water restriction for one week and maintained at ∼85% of their baseline body weight. Mice first underwent habituation to head fixation and reward delivery. After 2∼3 days of habituation, mice began shaping phase. Each trial began with a brief green light cue presented 3 seconds before odor onset, followed by the first odor, a 3-second inter-stimulus interval, a second odor presentation, and a 3-second response window. Odors were presented for either 2 seconds (Short) or 5 seconds (Long). During shaping, only nonmatch trials (Short-Long and Long-Short) were presented in a random sequence. Mice were trained to withhold licking during the first odor presentation and inter-stimulus interval and to lick during the response window to receive water reward. Once mice reached over 70% correctness on both trial types, they advanced to the full tDNMS task. During the tDNMS task, Short-Short trials were introduced in addition to the nonmatch trial types. For Short-Short trials, mice received no reward and were punished with an increased ITI (+12 s) for licking in the response window. The mice continued training until reaching the criterion (>70% correctness).

### Virtual Reality (VR) spatial task

The VR training began after animals reached criterion performance in the tDNMS task. The VR task was performed using the same behavioral apparatus as the tDNMS task. The virtual environment was implemented using a custom ViRMEn-based program^44,45^ in MATLAB. Visual stimuli were presented on five LCD monitors surrounding the head-fixed animal and were updated in real time according to locomotion on the treadmill. Mice were trained to navigate a 3-m linear virtual track containing multiple visual landmarks distributed along its length. Upon reaching the end of the track, animals received a water reward (∼5–6 μL), followed by a brief delay before being teleported back to the beginning of the track to initiate the next trial. Because mice were already familiar with head fixation and treadmill running from tDNMS training, they rapidly learned the VR task. Animals received at least four daily 30-min training sessions before recordings to ensure familiarity with the virtual environment and reward location.

### Dark session

Immediately following the tDNMS task, a 20-min Dark session was performed using the same apparatus as the tDNMS and VR tasks. During the Dark session, no rewards were delivered and no odor stimuli were presented. Locomotion was recorded through the same ViRMEn system used for VR experiments; however, all monitors displayed a black screen during Dark sessions. Mice spontaneously ran on the treadmill despite the absence of sensory cues or reward.

### Open field foraging task

Open field foraging task was conducted in a 30 x 40-inch (76 x 102 cm) rectangular arena with visual cues consisting of a striped pattern and a circular pattern attached to two walls of the arena. An overhead camera recorded the position of the animal during the task. To encourage exploration, small pieces of wet food pellet were scattered throughout the arena by the experimenter. Before recordings, mice were trained in the open field environment with at least four 20-min session. On recording days, open field foraging sessions were performed either before or after the tDNMS-Dark-VR recording sequence.

### Histology

Following completion of behavioral experiments, mice were perfused with 1x phosphate-buffered saline (PBS), followed by 4% paraformaldehyde (PFA). Brains were extracted, post-fixed in 4% PFA for approximately 24 h, and subsequently sectioned sagittally at 50-100 μm thickness using a vibratome. Sections were either stained with fluorescent NeuroTrace Nissl stain (Invitrogen) or mounted with a DAPI-containing mounting medium. Brain sections were imaged using a VS200 Virtual Slide fluorescence microscope (Olympus), and fluorescent probe tracks were used to verified Neuropixels recording locations. MEC boundaries were identified with reference to the Paxinos and Franklin mouse brain atlas based on cytoarchitectonic landmarks. The depth of each recorded unit was estimated from Kilosort 4 and registered to the histologically identified probe track to assign units to anatomical regions. Units located outside the MEC were excluded from subsequent analyses, and units from all MEC layers were pooled for analysis.

### Electrophysiology recording and preprocessing

Electrophysiological recording, preprocessing, and spike-sorting procedures were performed as previously described^27^. Briefly, electrophysiology data was recording at 30 kHz sampling rate using Neuropixels 2.0 probes and headstages (HS-2010, Imec) connected to the Imec acquisition system. Neural signals and behavioral signals from the tDNMS, Dark, VR, and open field foraging tasks were streamed into SpikeGLX software and synchronized. Because these behavioral tasks were performed sequentially within the same recording session, data from all tasks were concatenated and processed together, allowing individual units to be tracked across behavioral conditions.

Spike sorting was performed using Kilosort 4^46^ through the SpikeInterface framework^47^. Recording channels were grouped by shank and independently processed for high-pass filtering, phase-shift correction, common reference subtraction, removal of noisy channels, and motion correction using the SpikeInterface package. Following spike sorting, quality metrics were computed for all detected clusters using BombCell^48^. Custom BombCell parameters used for unit classification are available in the analysis code repository. Units classified as noise, multi-unit activity, or non-somatic units were excluded from further analyses.

### Data analysis

#### MEC principal cells

Following spike sorting and quality control using BombCell, putative principal cells were identified based on electrophysiological properties. Mean firing rate was calculated across the entire recording session. The peak-to-trough duration was computed based on the spike waveform extracted from the Kilosort templates. Units with a mean firing rate < 16 Hz and a peak-to-trough duration > 400 μs were classified as principal cells^49^. Only principal cells with a mean firing rate greater than 1 Hz were included in the following analyses.

#### Grid cell identification

Grid cells were identified based on distance-tuned activity during the Dark session, in which head-fixed mice ran freely on the treadmill in the absence of external sensory cues and reward. Firing rate maps were computed as a function of cumulative running distance using 3 cm bins. Spatial autocorrelograms were computed to visualize the periodicity of distance-dependent firing. Spatial periodicity was quantified by computing the power spectral density (PSD) of the rate maps using Welch’s method (scipy.signal.welch) with a Hann window (500-bin window) for frequencies between 0.2 and 2.5 cycles m^-1^. To determine statistical significance, spike trains were circularly shifted in a random offset 1,000 times, and PSD was recomputed for shuffled dataset. The neurons whose peak PSD exceeded the distribution of shuffled PSD by more than 5 standard deviations were classified as grid cells.

To validate the one-dimensional grid cell classification, the same neurons recorded during the open field foraging task were analyzed using conventional two-dimensional grid cell metrics. Grid cell metrics were calculated as previously described^50^. Two-dimensional firing rate maps were computed using 2 cm bins and smoothed with a Gaussian smoothing. Then two-dimensional spatial autocorrelograms were then computed from the firing rate map, and grid scores were calculated as the minimum rotational correlation at 60° and 120° minus the maximum correlation at 30°, 90°, and 150°. Grid spacing was estimated as the distance from the center of the autocorrelogram to the first ring of correlation peaks. To assess statistical significance, spike trains were circularly shifted relative to the behavioral trajectory by a random offset 1,000 times, and grid scores were recomputed for each shuffled dataset. Neurons with a grid score p < 0.05 and a grid spacing < 70 cm were classified as two-dimensional grid cells. Grid spacings exceeding 70 cm were excluded because the dimensions of the foraging chamber (76 × 102 cm) limited the reliable estimation of larger grid spacings.

#### Spatial cell identification

Spatial tuning during the VR task was quantified using spatial mutual information (MI). Position on the VR track was binned at 3 cm, and average firing rate maps across laps were computed along the track. The mutual information was calculated as follows^51^:

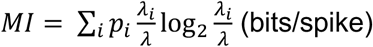

where *p_i_* is occupancy probability of spatial bin *i*, *λ_i_* is the mean firing rate in bin *i*, and *λ* is the overall mean firing rate. Statistical significance was assessed using 1,000 circular shift shuffles of spike trains, and p values were computed from the fraction of MI values of shuffled data greater than the observed MI. Neurons exhibiting significant spatial tuning (MI > 0.3 and p < 0.001) but not classified as grid cells were classified as non-grid spatial cells. Principal cells that met neither the grid cell nor spatial cell criteria were classified as non-spatial cells.

#### Time cell identification

Time cell identification was performed using correct trials from the tDNMS task. Neurons were classified as time cells if they met the following three criteria: 1) a significant time field, 2) high trial-by-trial temporal reliability, and 3) sufficient activity within the detected field. For time field detection, spike trains were binned into 250 ms time bins over a 10-s analysis window beginning at the onset of the first odor. Trial-by-trial firing rate maps were smoothed using a Gaussian rolling window (window size = 8 bins; standard deviation = 2 bins). For each trial type, spike trains from individual trials were circularly shifted 1,000 times. The candidate time field bins were defined as those whose firing rate exceeded the shuffle mean by more than five standard deviations. Three or more consecutive suprathreshold bins (750 ms) were grouped into a significant time field. Temporal reliability was quantified separately for each trial type using a leave-one-out trial-by-trial Pearson correlation analysis. For each trial, the Pearson correlation coefficient was calculated between the trial’s rate map and the average rate map of all remaining trials, and the correlation coefficients were averaged across trials. Statistical significance was determined by comparing the observed correlation coefficient with a shuffle distribution generated from 1,000 circular shifts within each trial. Trial participation was calculated as the fraction of trials in which the mean firing rate within the dominant time field exceeded the mean firing rate outside all detected fields. Neurons were classified as time cells if they contained at least one significant time field and satisfied all of the following criteria in at least one trial type: trial-by-trial correlation coefficient > 0.15, shuffle-derived p < 0.01, and trial participation > 40%.

#### Generalized linear model analysis

Generalized linear models (GLMs) were used to determine which coding scheme (spatial or temporal) is dominant during the VR and tDNMS tasks. Spike counts were binned at 100 ms time bins and modeled using Poisson regression implemented with the Python package statsmodels. Spatial and temporal predictors were generated using cubic splines with 10 degrees of freedom.

For the VR task, the spatial model used the animal’s position along the linear track as the predictor, and the temporal model used elapsed time within each lap. For the tDNMS task, the spatial model used cumulative distance traveled from trial onset, and the temporal model used elapsed time from trial onset. The null model assumed a constant firing rate equal to the mean firing rate across all position and time. Model performance was quantified using cross-validated log-likelihood obtained with 10-fold cross-validation across laps or trials. The performance of each model was expressed as the log-likelihood improvement relative to the null model

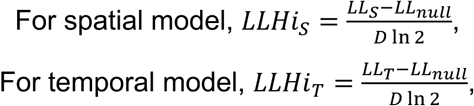

where *D* is the total data duration, and *LL_S_*, *LL_T_*, and *LL_null_* are log-likelihood of spatial, temporal, and null model. Positive *LLHi* values indicate that the corresponding model explained spikes better than the null model. To directly compare spatial and temporal coding within individual neurons, we calculated Δ*LLHi*:

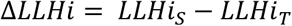

Positive Δ*LLHi* values indicate stronger spatial coding, and negative values indicate stronger temporal coding.

#### Rastermap

Grid cell population activity was sorted using Rastermap^52^ algorithms, which orders neurons according to the similarity of their activity patterns to visualize structured population activity. Long firing rate maps were generated for each neuron as a function of traveled distance with bin size 6 cm and normalized by z-scoring. Rastermap was performed using parameters as follows: n_clusters=None, n_PCs=32, locality=0.3, time_lag_window=20.

#### Time lagged cross correlation

Time-lagged cross-correlation analysis was adapted from the method previously described^28^. Neural firing rates were binned at 100 ms intervals and smoothed prior to analysis. A time-lagged cross correlation matrix was constructed to capture functional similarity between neurons while allowing for temporal offsets in their activity. For each neuron, its firing trace was shifted forward across a range of time lags (up to 30 time points), and each shifted version was compared with the firing-rate traces of all other neurons using a dot product. This produces, for every ordered neuron pair, a cross-correlation profile describing how strongly one neuron’s activity aligns with another neuron’s activity at different time delays. To make the measure symmetric between two neurons, the cross-correlation profile from neuron *_i_* to neuron *_j_* was combined with the reverse profile from neuron *j* to neuron *i*. The minimum-to-maximum ratio of the combined profile was then computed as the pairwise similarity score. Because this score becomes smaller when the combined profile contains a pronounced peak relative to its minimum, neuron pairs with lower min/max ratios were considered to exhibit stronger lag-dependent temporal relationships. These pairwise scores form a symmetric correlation-like matrix. Because the goal is to cluster neurons by similarity of their temporal interaction patterns, this matrix is then converted into a distance matrix using correlation distance applied to the squared similarity profiles. The resulting distance matrix reflects how similarly each neuron relates to the rest of the population across time lags. Finally, hierarchical agglomerative clustering is applied to this precomputed distance matrix, grouping neurons whose time-lagged correlation structure is similar, and the reordered distance matrix is visualized to reveal cluster structure among the neurons.

#### Low-dimension embedding

Low-dimension embedding analysis was adapted from the method previously described^28^. We begin with a selected time interval and divide it into evenly spaced time points. For each cell, the spike times that occur within this time range are used to build a smoothed firing activity trace sampled at those time points. Each spike is treated as a Gaussian-shaped bump centered at the spike time, so its contribution is largest at time points near the spike and smaller at time points farther away. At each sampled time point, the contributions from all nearby spikes of that cell are added together. This process is repeated for all cells, producing a firing matrix in which each row represents one cell and each column contains the estimated firing activity of the population at a particular sampled time point.

The firing activity matrix initially contains the activity of all recorded cells across the selected time points. This full matrix is first used to compute the time-lagged cross-correlation matrix, which identifies groups of cells with similar lagged activity patterns. We then restrict our analysis to the largest cluster of correlated cells. After selecting these cells, we further filter the time points by requiring sufficient overall neural activity, measured as the summed firing activity across the selected cells, and sufficient mouse movement speed. This produces a reduced activity matrix whose rows correspond to active time points and whose columns correspond to correlated cells. Each row is then treated as a data point representing the population neural activity of the mouse at a given moment in time. PCA is applied to this point cloud to reduce its dimension to 8. We then perform farthest-point subsampling, followed by denoising based on k-nearest-neighbor membership, yielding the final point cloud. Finally, a three-dimensional UMAP embedding was generated for visualization.

To quantify whether trial events were clustered in the neural state space, reward-onset and trial-onset population states were projected into the PCA space, and their mean pairwise Euclidean distance was compared against a null distribution. For null distribution, an equal number of time points was then randomly sampled from all analyzed (velocity-filtered) time points, and the mean pairwise Euclidean distance between the sampled population states was calculated. The procedure was repeated 1,000 times to generate the null distribution. P-value was computed as the fraction of null distribution whose mean pairwise distance was less than the observed value.

#### Statistical analysis

All analyses were performed in Python using NumPy, SciPy, pandas, scikit-learn, statsmodels, and custom analysis scripts. Unless otherwise stated, data are presented as mean ± SEM. Two-group comparisons were performed using two-sided Wilcoxon signed-rank tests or Wilcoxon rank-sum tests, as appropriate. Pearson correlation coefficients were used to quantify linear relationships between continuous variables. Associations between categorical variables were evaluated using Fisher’s exact test, chi-square tests of independence, or McNemar’s test where appropriate. Statistical significance was defined as *p* < 0.05.

## Acknowledgments

This work was supported by the National Institute of Mental Health of the National Institutes of Health (DP2MH129958 to J.G.H.), the National Science Foundation CAREER Award (IOS-2145814 to J.G.H.), and the Basic Science Research Program through the National Research Foundation of Korea funded by the Ministry of Education (RS-2023-00242639 to H.W.L.). We thank Dhruv Meduri and Bei Wang for generously sharing analysis codes that we adapted for use in this study.

## Contributions

H.W.L. and J.G.H. designed experiments and wrote the manuscript. H.W.L. and J.C.B. collected the data. H.W.L. and J.G.H. analyzed and interpreted the data.

